# Dysregulation of Acid Ceramidase-mediated Sphingolipid Metabolism Contributes to Tumor Progression in Tuberous Sclerosis Complex

**DOI:** 10.1101/2022.09.25.509382

**Authors:** Aristotelis Astrinidis, Chenggang Li, Erik Y. Zhang, Xueheng Zhao, Shuyang Zhao, Minzhe Guo, Rong Huang, Alan G. Zhang, Elizabeth Kopras, Nishant Gupta, Eric Smith, Elizabeth Fugate, Diana Lindquist, Kathryn Wikenheiser-Brokamp, Kenneth D. Setchell, Francis x. McCormack, Yan Xu, Jane J. Yu

**Affiliations:** Department of Internal Medicine, University of Cincinnati College of Medicine; Cincinnati Children’s Hospital Medical Center

**Keywords:** tuberin (TSC2), acid ceramidase (ASAH1), sphingolipid metabolism, tumorigenesis, single cell RNA sequencing, mTOR inhibitor rapamycin, tuberous sclerosis complex, lymphangioleiomyomatosis

## Abstract

Tuberous Sclerosis Complex (TSC) is disorder of multi-system benign neoplasia in the brain, heart, kidneys and lungs. Lymphangioleiomyomatosis (LAM) is a progressive pulmonary disease affecting exclusively women. Both are caused by mutations in *TSC1* and *TSC2*, resulting in mTORC1 hyperactivation. Single cell RNA sequencing of LAM lungs identified activation of genes in the sphingolipid pathway. Independent validation studies showed that acid ceramidase (*ASAH1*) and dihydroceramide desaturase (*DEGS1*), key enzyme for regulating sphingolipid and ceramide metabolism, were significantly increased in TSC2-null cells, and their expression and activity were rapamycin-insensitive. TSC2 negatively regulated the biosynthesis of tumorigenic sphingolipids. Suppression of ASAH1 by shRNA or the inhibitor ARN14976 (17a) markedly decreased the viability of TSC2-null cells. *In vivo*, 17a significantly decreased the growth of Tsc2-null cell derived mouse xenografts. When combined with rapamycin, 17a more strongly inhibited the progression of renal cystadenomas in *Tsc2^+/-^* mice than either agent alone, evaluated by pathology and MRI. Collectively, our studies identify a rapamycin-insensitive disorder of sphingolipid metabolism in TSC2-null cells and tumors and validate the novel hypothesis that TSC2 regulates sphingolipid production and action via ASAH1. Targeting aberrant sphingolipid metabolism pathways may have therapeutic value in TSC and LAM, and possibly in mTORC1-hyperactive neoplasms.

## Introduction

Tuberous Sclerosis Complex (TSC) is an autosomal dominant disorder with multi-system manifestations including benign neoplasms in the brain, heart, lungs, and kidneys (1–3). Complications associated with TSC tumors include cognitive impairment, intractable seizures, autism, renal hemorrhage and insufficiency, and respiratory failure. TSC is caused by germline inactivating mutations in *TSC1* or *TSC2* (4–6), which encode hamartin (TSC1) and tuberin (TSC2). These form a protein complex with GTPase activating protein activity toward Ras homolog enriched in brain (Rheb), an activator of the mechanistic target of rapamycin complex 1 (mTORC1) (7–9). Thus, loss of either TSC1 or TSC2 results in constitutive activation of mTORC1. mTOR is a serine/threonine protein kinase that functions as the catalytic subunit in two complexes, mTORC1 and mTORC2 (10). mTORC1, containing Raptor, phosphorylates a plethora of substrates including ribosomal protein S6 kinase (S6K1/2), the translational repressor 4E-binding protein 1 (4E-BP1) and ULK1, to regulate ribosome biogenesis, mRNA translation, protein synthesis, autophagy, cell metabolism, gene transcription, and cell growth (11–13). mTORC2, containing Rictor, phosphorylates protein kinases including AKT at Serine 473 and PKC to control cell survival, cytoskeletal organization and cell motility (14). The understanding that *TSC1* or *TSC2* mutations cause dysregulation of mTORC1/mTORC2 led to preclinical studies (15–20) and human clinical trials demonstrating the effectiveness of mTORC1 inhibitors (21–26). However, the effect of mTORC1 inhibitors (mTORi) on tumor growth has been consistently cytostatic rather than cytotoxic; tumors typically regrow upon the cessation of treatment (21, 23, 27–29), some tumors become resistant on therapy and all patients are at risk of potential side effects from long-term treatment. Thus, there is a need for both improving the efficacy of mTORi and deriving strategies in which combination with other agents could induce curative cytotoxic effects.

Sphingolipids, major components of eukaryotic plasma membrane, play key roles as mediators of multiple cellular functions, including apoptosis, proliferation, stress responses, necrosis, inflammation, autophagy, senescence, tumor growth (30, 31), cancer progression, and differentiation (31). Ceramide and sphingosine-1-phosphate (S1P), the two central bioactive sphingolipids, exhibit opposing roles in regulating cancer cell death and survival (31–33). Ceramide is considered to be at the core of sphingolipid metabolism and intracellular concentrations of this intermediary directly increase apoptosis and cell cycle arrest (34). *De novo* synthesis of ceramide begins with the condensation of palmitate and serine, the rate-limiting step being catalyzed by serine palmitoyl transferase. Subsequently, the final reaction to produce ceramide from dihydrocramide is catalyzed by Delta 4-Desaturase Sphingolipid 1 (dihydroceramide desaturase, DEGS1). N-Acylsphingosine Amidohydrolase 1 (acid ceramidase, ASAH1) catalyzes the cleavage of ceramide to sphingosine and fatty acids, and controls the sphingosine/ceramide ratio (31, 34). Importantly, carmofur, a potent *in vivo* inhibitor of ASAH1 activity (35), has been used in the clinic to treat colorectal cancers in other countries (36), although not persued for FDA approval.

We previously published that prostaglandin biosynthesis and actions are dysregulated in TSC2-deficient cells (37–40). Prostaglandin E2 receptors (EP1-4) are G protein-coupled receptors that integrate signals to regulate various cellular functions (41, 42) such as cyclic AMP production and transactivation of gene expression including *ASAH1* (43). We have found that TSC2-null angiomyolipoma-derived cells produce excessive amount of PGE_2_ via COX2 and express higher levels of PGE_2_ receptor 3 (EP3) relative to TSC2-reexpressing cells (37–40). These results suggested an interplay of PGE_2_ action and sphingolipid metabolism in TSC tumors. In the present study, we analyzed publicly available expression array data from TSC2-null angiomyolipoma-derived cells (44) and found that the transcript levels of *DEGS1* and *ASAH1* were significantly increased in TSC2-null angiomyolipoma-derived cells. In addition, we discovered TSC2-deficiency dependent upregulation of *DEGS1* and *ASAH1* expression in cell models of human angiomyolipoma-derived cells and in clinical specimens of renal angiomyolipomas and pulmonary LAM nodules. Mechanistically, enhanced expression of DEGS1 and ASAH1 is independent of mTORC1 inhibition, which is novel and previously unexplored. Together, our studies identify a rapamycin-insensitive sphingolipid metabolism in TSC2-null cells and tumors. Our studies reveal a novel function of TSC2 that negatively regulates sphingolipid production and action via ASAH1. Thus, targeting aberrant sphingolipid metabolism pathways may have therapeutic values in TSC and LAM, and possibly in mTORC1-hyperactive neoplasms.

## Results

### Single cell RNA-sequencing identifies activation of sphingolipid metabolism pathway genes in LAM lung lesion cells

We utilized scRNAseq and systems biology tools to identify LAM cells, characterize cells based on their signature genes and enriched functions, and to delineate altered signaling pathways associated with LAM pathogenesis. Data mining of LAM lung (n=12,374 cells) identified a LAM^core^ cluster (**Fig. 1a**), which selectively express known LAM signature genes *(PMEL, ACTA2, FIGF, and GPNMB)* (**Fig. 1b**). Functional enrichment analysis and IPA analysis revealed aberrant activation of sphingolipid signaling pathway (**Fig. 1c**) and associated genes, including *S1PR3*, induced in LAM^core^ but not in mesenchyme, endothelial, epithelial, or immune cells (**Fig. 1d).**

**Figure 1.**
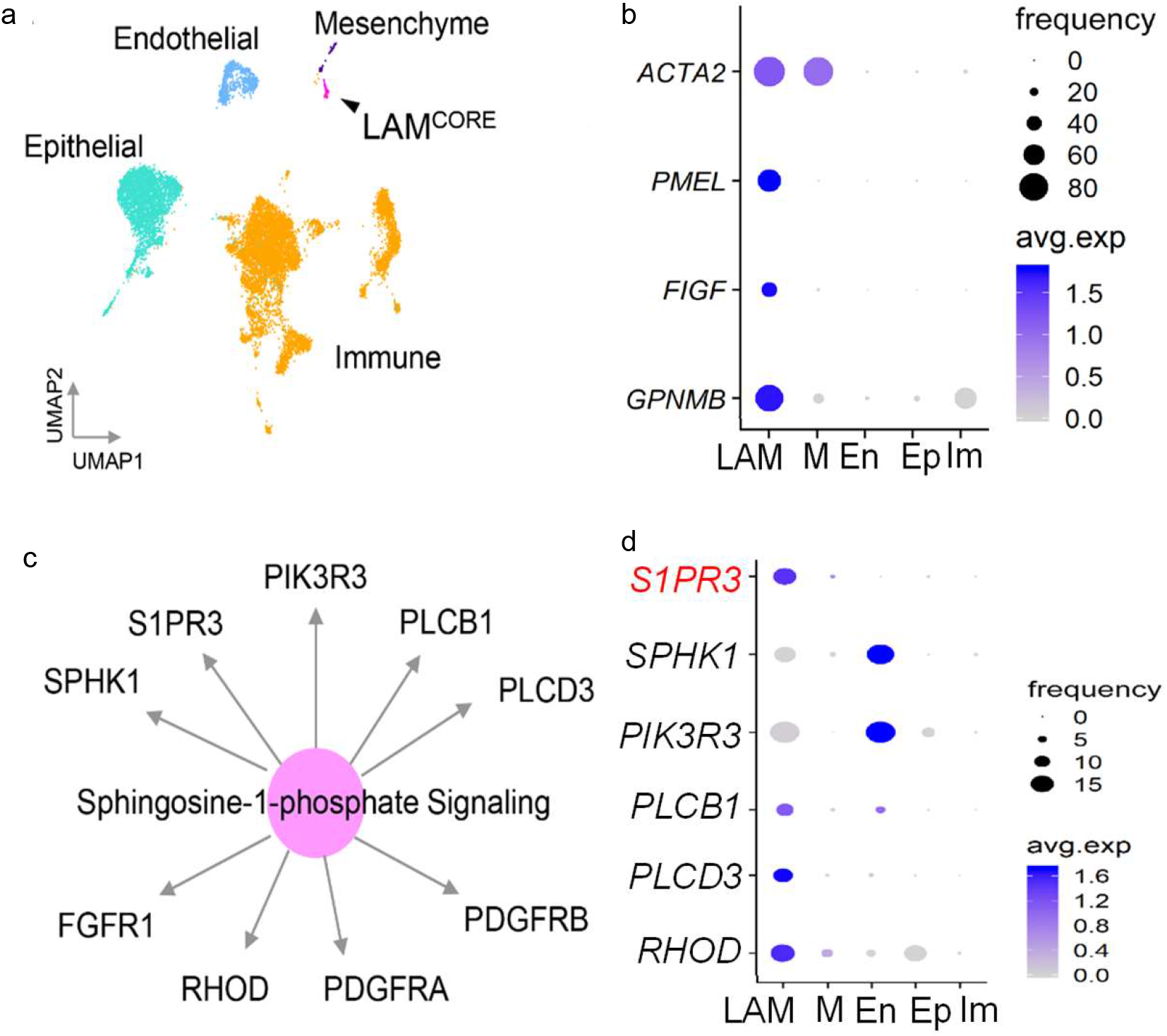
LAM single cell RNAseq analysis identifies activation of sphingolipid pathway associated genes in LAM^core^ cells. (a) scRNAseq analysis of LAM lung (n=12,374 cells) identifies a LAM^core^ cell population, which selectively expresses known LAM-related genes (b). (c) Sphingolipid pathway and associated genes were induced in LAM^core^ cells. (d) Expression of representative genes including S1PR3. Node size is proportional to the expression frequency of a gene in a cell population. Blue color indicates high expression. LAM, LAM^Core^; M, Mesenchyme; En, Endothelial; Ep, Epithelial; Im, Immune cells.

### TSC2 negatively regulates the expression and activity of ASAH1

To determine whether sphingolipid metabolism is regulated by TSC2, we re-analyzed previously published expression array data from TSC2-null angiomyolipoma-derived cells (44). We found that the transcript levels of *DEGS1* and *ASAH1* were significantly increased in TSC2-null cells, compared to TSC2-addback cells (**Fig. 2a** and **2b**). Immunoblotting analysis showed that the protein levels of DEGS1 and ASAH1 were markedly higher in TSC2-null cells relative to TSC2-addback cells, both in regular (10% FBS) and serum-free (0% FBS) culture conditions (Fig. **2c** and **2d**). These findings indicate aberrant upregulation of the sphingolipid metabolism pathway in TSC2-null angiomyolipoma-derived cells.

**Figure 2.**
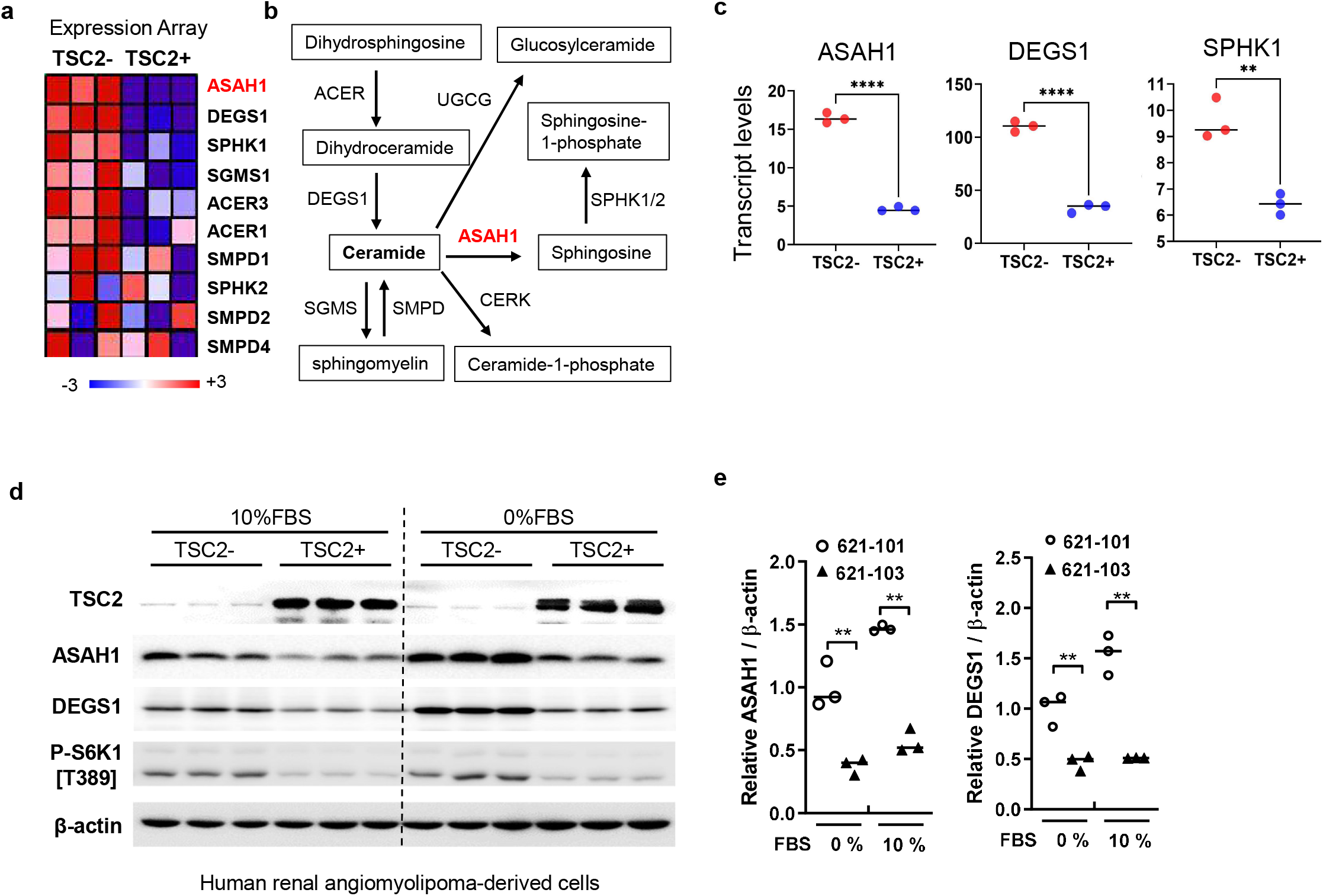
Upregulation of dihydroceramide desaturase (DEGS1) and acid ceramidase (ASAH1) expression in TSC2-null LAM-derived cells. (a) A heatmap of the expression of sphingolipid metabolism pathway genes in TSC2-null (TSC2-) 621-102 and TSC2-addback (TSC2+) 621-103 cells. The scale indicates the fold change of genes from blue (min) to red Max (−3 to +3). (b) Sphingolipid metabolism pathway shows upregulated expression of genes in the pathway. (c) Transcript levels of *ASAH1, DEGS1* and *SPHK1* in TSC2-null patient-derived 621-101 cells. (d) The protein levels of TSC2, DEGS1, ASAH1, and phospho-S6K1 (P-S6K1) (T389) were assessed by immunoblotting. β-actin was used as a loading control. (e) Densitometry of ASAH1 and DEGS1 protein levels normalized to β-actin (n=3/group). **p<0.01, Student’s t-test.

### Differential expression of acid ceramidase ASAH1 is evident in Tuberous Sclerosis Complex lesions

Activation of the sphingolipid metabolism pathway leads to sphingosine production and activation of bioactive lipid signaling molecules, thereby increased cell viability and tumor progression (31, 34). Note that angiomyolipomas have “mesenchymal” characteristics, containing immature smooth cells, fat cells, and aberrant blood vessels (45). To determine whether the sphingolipid metabolism pathway is activated in human TSC2-null lung lesions, we extracted gene expression array data from LAM lungs (n=14) and non-LAM female lungs (n=15), and found that *DEGS1* and *ASHA1* are upregulated in LAM tissues, but not in non-LAM female lung cells (**Fig. 3a and 3b**). To determine the protein levels of ASAH1 in human lung lesions we performed immunohistochemical staining in LAM lung tissues obtained from the NDRI. Expression of ASAH1 coincided with smooth muscle actin positive and phospho-S6 positive LAM lung lesions, compared to adjacent normal tissue (**Fig. 3c**). Finally, we assessed the protein levels of ASAH1 in LAM lung vs. normal lung tissue lysates (**Fig. 3d**) and found that ASAH1 levels in LAM lungs were increased by approximately 4-fold, compared to normal lung (**Fig. 3e**). Since ASAH expression is regulated by cAMP-responsive element binding protein (CREB) (46), we tested the activation of CREB by immunoblotting with a phospho-CREB [S133] antibody and found evidence of a two-fold increase of the p-CREB/CREB ratio (**Fig. 3d and 3e**). Collectively, these data suggest activation of the sphingolipid biosynthesis pathway in TSC2-null cells and lesions.

**Figure 3.**
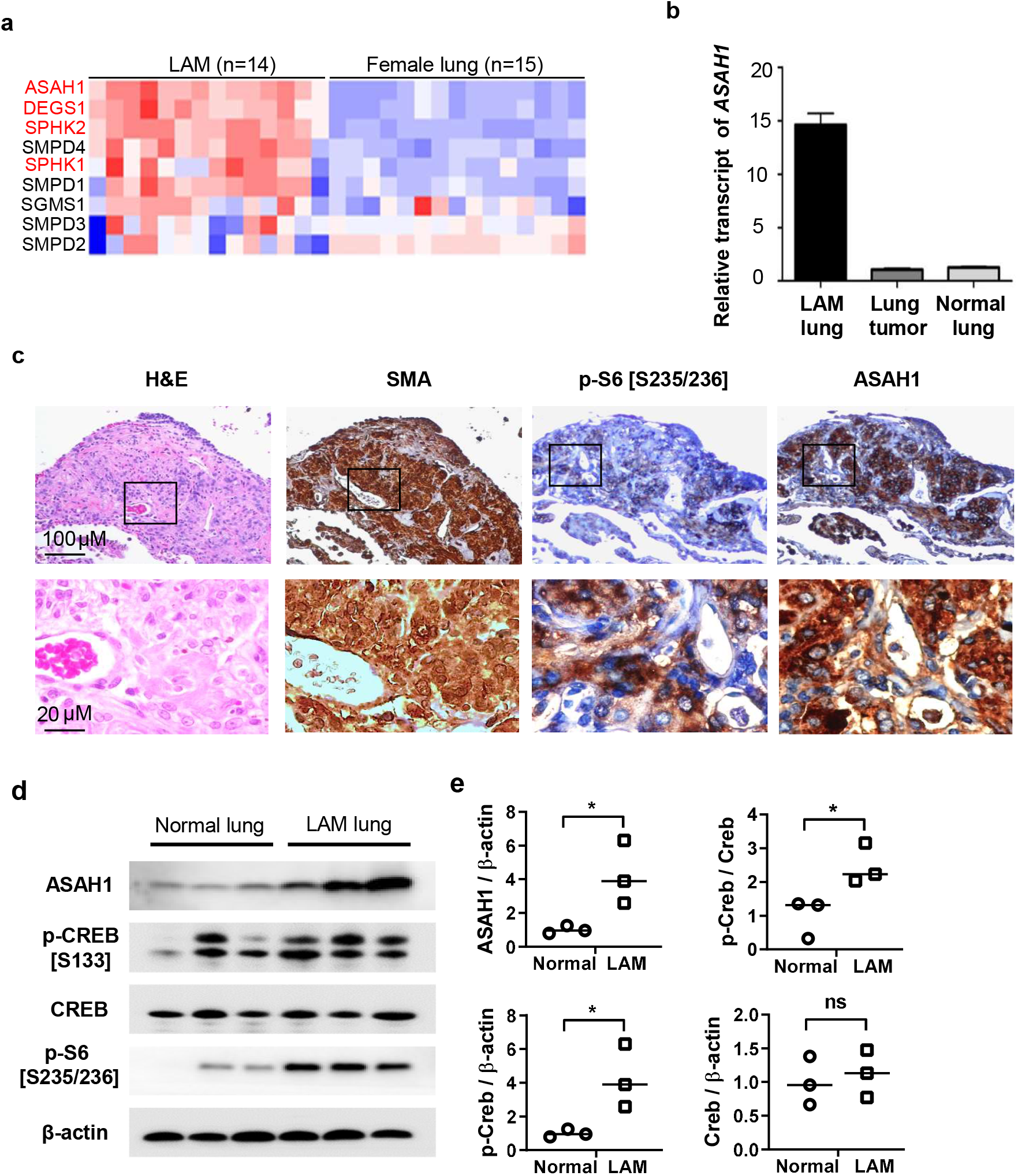
Expression of DEGS1 and ASAH1 is evident in pulmonary LAM lesions. (a) Heatmap of gene expression of ASAH1 and DEGS1 in laser capture microdissected LAM lesion cells (n=14 subjects) and non-LAM female lungs (n=15 subjects). (b) Realtime RT-PCR analysis of *ASAH1* in LAM lung and control female lungs. (c) Immunohistochemistry of hematoxylin and eosin (H&E), smooth muscle actin (ACTA2), phospho-S6 (S235/236), and ASAH1 in LAM lung tissues. Representative images are shown. (d) Immunoblotting of ASAH1 and transcription factor CREB in LAM lung and control lung (n=3). (e) Densitometry of ASAH1 and phospho-CREB (n=3). *p<0.05, Student’s t-test.

### ASAH1 activity is elevated in a rapamycin-insensitive manner in LAM-derived cells

ASAH1 catalyzes the cleavage of ceramide to sphingosine. To determine whether the elevated ASAH1 expression in TSC2-null cells is correlated with its enzymatic activity, we measured the cellular levels of ceramides and sphingosine using LC-MS/MS. In TSC2-null cells the levels of ceramides were three-fold lower (**Fig. 4a**), and the level of sphingosine was two-fold higher (**Fig. 4b**), compared to TSC2-addback cells. Importantly, rapamycin treatment did not alter the levels of ceramides or sphingosine in TSC2-null cells, suggesting a rapamycin-insensitive regulation of ASAH1 activity. Moreover, qRT-PCR analysis showed that ASAH1 mRNA levels were 2.4-fold higher in TSC2-null cells. compared to TSC2-addback cells, and rapamycin did not affect ASAH1 mRNA levels (**Fig. 4c**). Furthermore, ASAH1 protein levels were higher in TSC2-null cells, compared to TSC2-addback cells, and rapamycin did not alter ASAH1 levels while suppressing S6 phosphorylation as expected (**Fig. 4d**). We next treated TSC2-null cells with rapalink-1 (RLK1, 5 nM), a third generation mTOR inhibitor (mTORi) that is more potent than rapamycin and other first- and second-generation mTORi. Although rapalink-1 treatment drastically suppressed phosphorylation of S6 and 4E-BP1, ASAH1 protein level was not affected (**Fig. 4e**). Collectively, our data indicate an mTOR-independent regulation of ASAH1 expression and activity, and sphingolipid metabolism in angiomyolipoma-derived TSC2-null cells.

**Figure 4.**
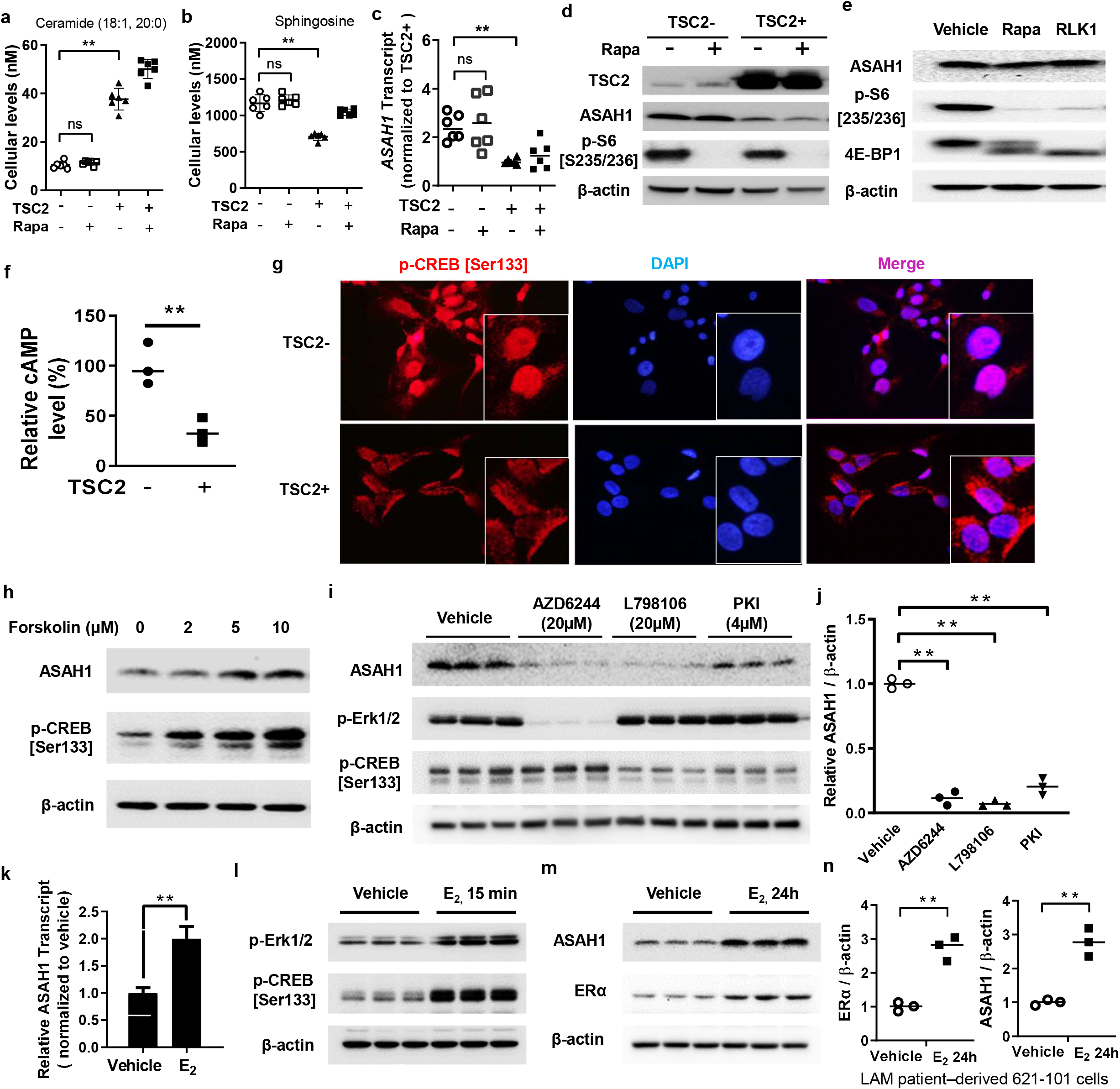
ASAH1 activity and expression is elevated in a sirolimus-insensitive manner in LAM-derived cells. 621-101 (TSC2-) and 621-103 (TSC2+) cells were treated with 20 nM sirolimus (Rapa) for 24 hr. Cellular levels of ceramide (a) and sphingosine (b) were quantified using LC-MS/MS. (c) *ASAH1* transcript levels were quantified by RT-PCR. (d, e) LAM patient-derived 621-101 (TSC2-) and 621-103 (TSC2+) cells were treated with mTORC1 inhibitor rapamycin (20 nM) or rapalink-1 (RLK1) (0.1 μM) for 24 hr. Protein levels of TSC2, ASAH1, phospho-S6 (S235/236), and 4E-BP1 were assessed by immunoblotting. β-actin was used as a loading control. (f) Cellular levels of cAMP were quantified in 621-101 (TSC2-) and 621-103 (TSC2+) cells (n=3). (g) Representative images of confocal microscopy of phospho-CREB were illustrated. Nuclei were stained with DAPI. (h, i) 621-101 (TSC2-) cells were treated with cAMP agonist forskolin (2, 5, and 10 μM), MEK1/2 inhibitor AZD6244 (20 μM), prostaglandin E2 receptor 3 (EP3) inhibitor L798106 (20 μM), or protein kinase A inhibitor (PKI) (4 μM), for 24 hr. Protein levels of ASAH1, phospho-CREB (S133), and phospho-Erk1/2 were assessed by immunoblotting. β-actin was used as a loading control. (j) Densitometry of ASAH1 protein levels normalized to β-actin (n=3). (k-m) 621-101 (TSC2-) cells were treated with 10 nM E_2_ for 15 min or 24 hr. *ASAH1* transcript levels were quantified using RT-PCR. Immunoblot analyses of phospho-CREB (S133), and phospho-Erk1/2, ASAH1 and ERα were performed (n=3/group). β-actin was used as a loading control. (m) Densitometry of ERα and ASAH1 protein levels (n=3). **p<0.01, Student’s t-test.

### ASAH1 expression is dependent on CREB and ERK activity

It was previously reported that cAMP inhibits mTORC1 signaling in a TSC1/TSC2-independent manner (47). cAMP response element-binding protein (CREB) is tightly regulated by intracellular cAMP levels, and since we observed an upregulation of CREB phosphorylation in LAM lung tissue lysates (Fig. 3d), we assessed the cAMP levels in TSC2-null and TSC2-reexpressing cells and found that expression of TSC2 decreased intracellular cAMP levels by 3-4-fold (**Fig. 4f**). Consistent with the observations, TSC2-null cells had increased phospho-CREB (Ser133) staining, compared to TSC2-reexpressing cells (**Fig. 4g**). To test whether increased CREB activation correlates with ASAH1 expression, we treated TSC2-null cells with forskolin, an adenyl cyclase activator leading to increased intracellular cAMP. ASAH1 protein levels were increased in a dose-dependent manner (**Fig. 4h**). As expected, CREB phosphorylation was also increased by forskolin. Since activation of various pathways increase the phosphorylation of CREB, including MAPK, Protein Kinase A (PKA), prostaglandins, and estrogen (48), we tested whether perturbations of these pathways lead to alterations in ASAH1 levels in TSC2-null cells. Inhibition of MEK1/2, prostaglandin E2 receptor 3 (EP3), or PKA all lead to a significant decrease of ASAH1 protein levels (**Fig. 4i** and **4j**) and, except for MEK1/2 inhibition, to a decrease in CREB phosphorylation (**Fig. 4i**). Finally, estradiol (E2) led to increased transcript levels of ASAH1 by 2-fold (**Fig. 4k**), an immediate phosphorylation of CREB within 15 minutes (**Fig. 4l**), and increased ASAH1 protein levels by 2.7-fold concomitant with 2.8-fold increase of ERα within 24 hours (**Fig. 4m** and **4n**). Collectively, these data suggest that ASAH1 transcription and protein levels are dependent on CREB transcriptional activity.

### ASAH1 inhibitors selectively reduce the viability of TSC2-null cells

To determine the effect of ASAH1 blockade on cell viability, we treated angiomyolipoma-derived 621-101 cells with the ASAH1 inhibitor 17a (49). 17a selectively decreased the viability of TSC2-null cells in a dose-dependent manner with an IC_50_=117 nM (**Fig. 5a**), but not of TSC2-addback 621-103 cells (IC_50_=2,560 nM). Carmofur is a potent ASAH1 inhibitor that has been used to treat colorectal cancer patients in other countries (35, 50). Carmofur treatment also selectively reduced the viability of TSC2-null cells in a dose-dependent manner (**Fig. 5b**, IC_50_=17 μM), and to a lesser extent of TSC2-addback cells (IC50= 253 μM). These data support an important role of ASAH1 in TSC tumor cell viability.

**Figure 5.**
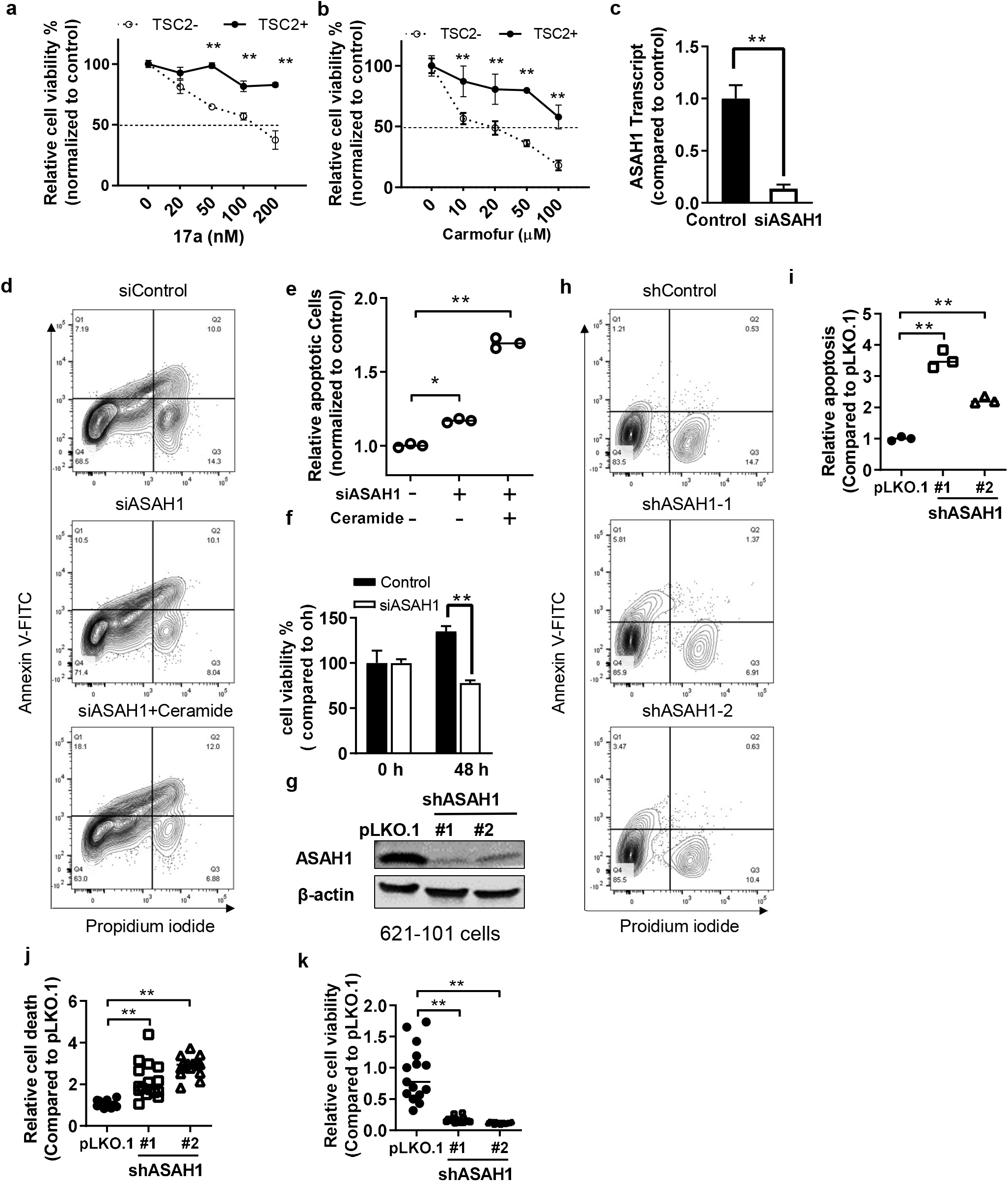
Suppression of ASAH1 decreases the survival of TSC2-null cells in vitro. 621-101 (TSC2-) and 621-103 (TSC2+) cells were treated with an ASAH1 inhibitor 17a (a) or carmofur (b) with indicated concentrations for 72 hr. Cell viability was measured using MTT assay (n=6/treatment group). (c) TSC2-null 621-101 cells were transfected with three independent ASAH1-siRNAs or control-siRNA for 48 hr. siRNA knockdown efficiency was determined by RT-PCR (n=3/group). (d) Cells were treated with ceramide, and then stained with Annexin V: FITC Apoptosis Detection Kit (BD#556547). Cell death was analyzed by flow cytometry (n=3). (e) The percentage of apoptotic (Annexin V+) cells was determined (percentage of Annexin V: FITC-positive cells in total cell number). (f) Cell viability was assessed 48 hr post siASAH1 RNA transfection in 621-101 cells using MTT assay. (g) siRNA knockdown efficiency was determined by immunoblotting. β-actin was used as a loading control. (h) 621-101 cells were infected with lentiviruses containing shRNA for vector pLKO, or *ASAH1*, and then selected with puromycin for two weeks. Stable cells were harvested, and then stained with Annexin V: FITC Apoptosis Detection Kit. Cell death was analyzed by flow cytometry (n=3). (i) The percentage of apoptotic (Annexin V+) cells was determined (percentage of Annexin V: FITC-positive cells in total cell number). (j) Cell death was measured using PI exclusion assay. Relative cell death was compared between pLOK.1 and shRNA-*ASAH1* cells. (k) Cell viability was measured using MTT assay (n=8-16/treatment group). *p<0.05; **p<0.01, Student t-test.

### Molecular depletion of ASAH1 inhibits the growth and induces apoptosis in angiomyolipoma-derived cells

To determine whether ASAH1 is a key mediator of TSC2-null cells growth we depleted ASAH1 using siRNA. 621-101 cells transfected with ASAH1 siRNA had 86.4% reduction of *ASAH1* transcript levels measured by qRT-PCR (**Fig. 5c**). ASAH1 siRNA-transfected 621-101 cells exhibited 42% reduction of cell viability compared to control siRNA (**Fig. 5d**), further supporting a critical role of ASAH1 in TSC tumor cell viability. More importantly, ASAH1 siRNA-transfected cells showed a significant increase in apoptosis compared to control siRNA transfected cells (**Fig. 5e** and **5f**), which was further increased by ceramide treatment. Finally, we transduced TSC2-null 621-101 angiomyolipoma-derived cells with lentiviruses having the pLKO.1 control vector or the ASAH1 shRNA (**Fig. 5g**). Consistent with our previous results, ASAH1 silencing resulted in a significant inhibition of cell viability (**Fig. 5h**) and a significant increase in apoptosis (**Fig. 5i-k**).

### Suppression of ASAH1 attenuates the progression of xenograft tumors of TSC2-null cells

We next assessed the possible benefit of ASAH1 inhibitor 17a in a xenograft tumor model of Tsc2-null ELT3 luciferase-expressing cells (20). Upon tumor development mice were treated with 17a or vehicle control for four weeks. Bioluminescence imaging was performed weekly. 17a treatment decreased the intensity of bioluminescence **(Fig. 6a)** and slowed down tumor progression by 2.7-fold, relative to the vehicle control (**Fig. 6b**). To further assess the role of ASAH1 in tumor growth, angiomyolipoma-derived 621-101 luciferase-expressing cells were transfected with shRNA-ASAH1 (**Fig. 6c**) and inoculated in NSG mice. Bioluminescence signal was detectable at 21 weeks post inoculation (**Fig. 6d**), and tumor growth was monitored until week 32 (**Fig. 6e**). ASAH1 depletion significantly delayed the onset of tumors and decreased tumor progression by 4.2-fold compared to pLKO.1 control transduced cells (**Fig. 6f**). Finally, we assessed whether suppression of ASAH1 affects the ability of TSC2-null cells to colonize the lungs in a mouse model similar to the one we previously reported (20). Female NSG mice were pre-treated with 17a or vehicle for 48 hours prior to intravenous inoculation with luciferase-expressing 621-101 cells, and bioluminescence imaging in the thoracic region was used to quantify lung colonization at baseline and at 6 and 24 hours post injection (**Fig. 6g**). Inhibition of ASAH1 by 17a significantly inhibited 621-101-induced lung colonization at both timepoints (**Fig. 6h**). Similarly, we inoculated NSG mice with the same number of 621-101 luciferase-expressing cells transduced with pLKO.1 control shRNA or with two ASAH1 shRNAs and performed bioluminescence imaging at baseline and 6 and 24 hours post inoculation (**Fig. 6i**). At both timepoints, both ASAH1 shRNAs significantly inhibited 621-101 cell lung colonization, compared to pLKO.1 shRNA control cells (**Fig. 6j)**. Collectively, these data support the notion that ASAH1 contributes to TSC tumor progression and dissemination to the lungs in LAM. We postulate that combinatorial suppression of mTORC1 and ASAH1 will potently suppress tumor growth.

**Figure 6.**
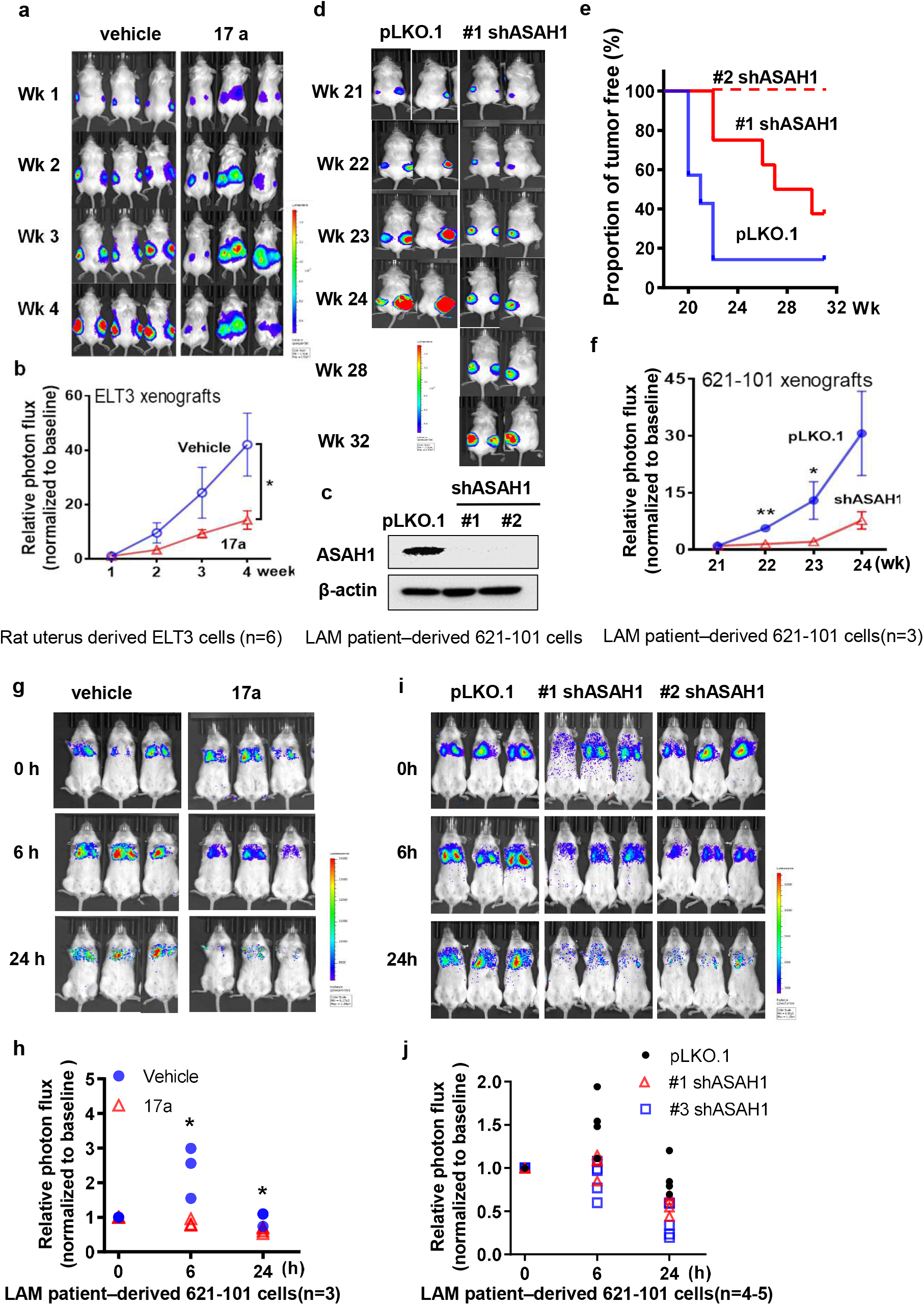
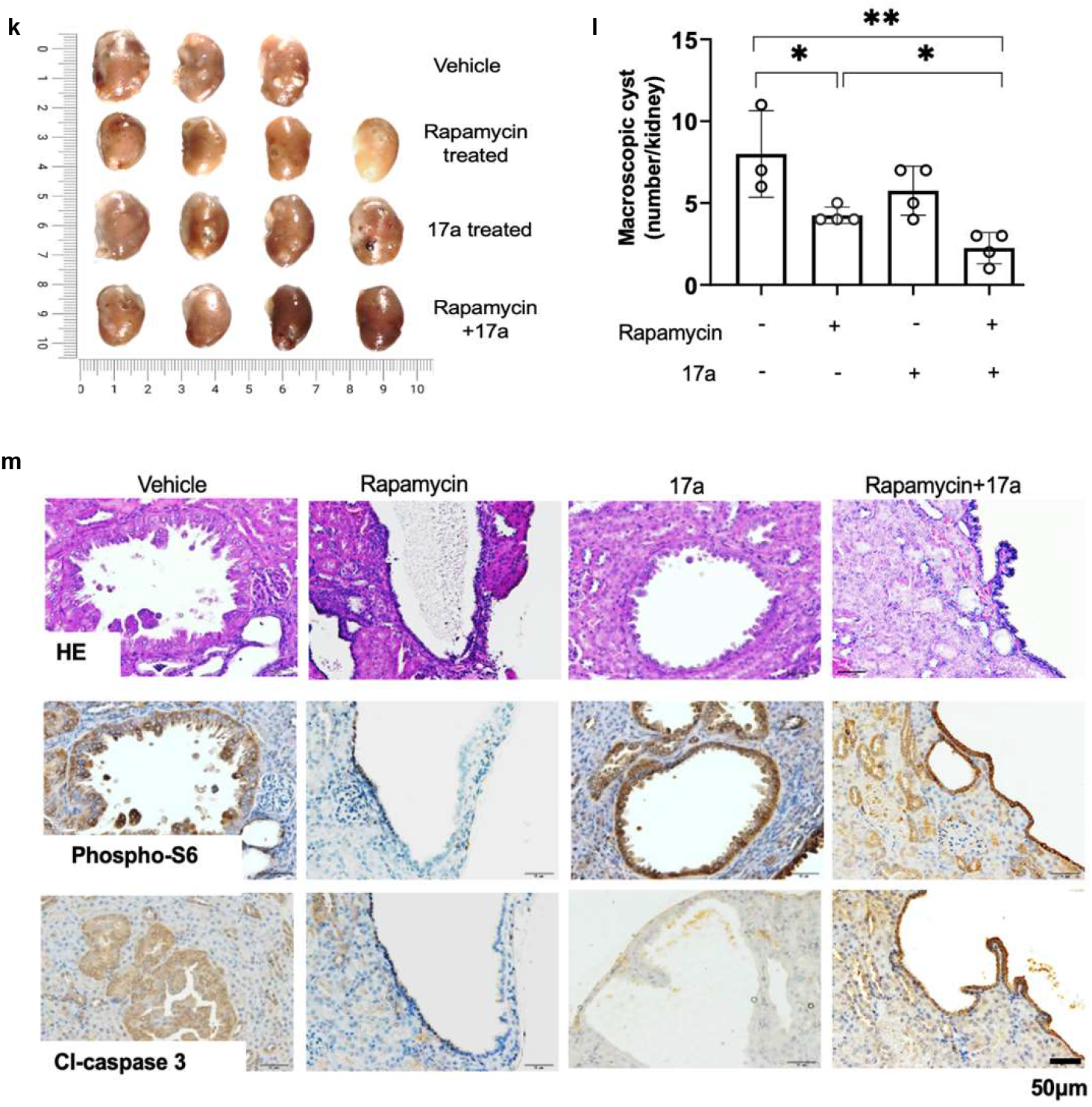
Suppression of ASAH1 decreases the survival of TSC2-null cells in vivo. (a) Female NSG mice were inoculated subcutaneously with 2×10 ELT3-luciferase expressing cells. Weekly bioluminescent imaging was performed. Upon ELT3 tumor onset, mice were randomized and treated with vehicle or 17a (10 mg/kg/day, i.p.) for four weeks. (b) Tumor photon flux was quantified and normalized to the baseline measurements (Week 0). Weekly bioluminescent imaging was performed. (c) 621-101 cells expressing luciferase (621L9) cells were infected with lentivirus of ASAH1-shRNA cells or control pLKO.1 empty vector. *ASAH1* knockdown efficiency was determined by immunoblotting. β-actin was used as a loading control. (d) Female NSG mice were subcutaneously inoculated with 2×10 621-101-pLKO.1 or 621-101-shASAH1 cells. Tumor formation was detected 21-weeks post cell inoculation. Weekly bioluminescent imaging was performed up to 32 weeks. (e) Kaplan-Meier percent tumor-free survival curve. (f) Tumor photon flux was quantified and normalized to the baseline measurements (Week 21) (n=3-4/group). (g) Female NSG mice were treated with vehicle or 17a (10 mg/kg/day, i.p.) for two days, and then intravenously inoculated with 2 x 10 621L9 cells. Bioluminescent imaging was performed 1-24 hours post cell inoculation (n=3). (h) Photon flux at the chest region was quantified. (i) Female NSG mice were intravenously inoculated with 2 x 10 621L9-ASAH1-shRNA (#1 and #2) cells or control pLKO.1 cells. Bioluminescent imaging was performed 1-24 hours post cell inoculation (n=4-5). (j) Photon flux at the chest region was quantified. *p<0.05, **p<0.01, Student t-test. (k) Tsc2+/- A/J mice were given vehicle, rapamycin, 17a, rapamycin and 17a combination treatment for 12 weeks (n=4 mice/group). Macroscopic analysis and (l) quantification of renal tumor burden under dissection microscope upon drug withdrawal. *p<0.05, **p<0.01, Student t-test. (m) Histology analysis of renal tumor burden upon drug withdrawal. H&E, immunohistochemistry staining of phospho-S6 and cl-Caspase 3 were performed.

### Acid ceramidases 1 inhibitor 17a in combination with rapamycin suppresses development and progression of renal cystadenomas in Tsc2 heterozygous mice

We have demonstrated the therapeutic benefits of the combinatorial treatment of 17a and rapamycin in intravenous and subcutaneous models. Next, we used Tsc2 heterozygote mice that develop spontaneously arising renal cystadenomas. Numbers of macroscopic renal lesions were decreased significantly in mice treated with 17a or rapamycin at week 12 (**Fig. 6k**). Importantly, the combinatorial treatment of 17 and rapamycin potently reduced the number of renal cystadenomas (**Fig. 6k and 6i**). Moreover, histological analysis showed that the dual treatment led to robust reduction of cell proliferative marker PCNA and increase of cell apoptotic marker cleaved caspase-3 from renal tumors compared to that from either single treatment (**Fig. 6m**). Collectively, our results demonstrate that inhibition of ASAH1 and mTORC1 potently suppresses tumor growth in preclinical models.

## Discussion

Sphingolipids play critical roles in kidney physiology (52), chronic inflammation and cancer progression (41, 42). Various sphingolipids can have widely differing cellular actions that likely contribute to cell viability in renal angiomyolipoma-derived cells. We have shown aberrant expression of DEGS1 and ASAH1, and change in production of sphingolipids in renal angiomyolipoma-derived cells. We also found that ASAH1 suppression led to reduced viability and increased apoptosis of TSC2-null cells. Furthermore, TSC2 negatively regulates the expression and activity of ASAH1 in an mTOR inhibitor-independent manner. Importantly, renal angiomyolipomas accumulate abundant levels of DEGS1 and ASAH1.

Our data shows that rapamycin, while successfully suppressing mTORC1 activation as assessed by phospho-S6 levels, had no effect on ASAH1 expression or activity in TSC2-null cells. However, studies by Choo et al. have shown that rapamycin’s inhibition of mTORC1 is incomplete (53). Moreover, mTORC2 dysfunction has been observed in TSC2-null cells (54), and there is further evidence that TSC2 may have other functions in addition to inhibiting mTORC1 or mTORC2. Here we propose to determine the mechanism by which TSC2 regulates the expression of sphingolipid biosynthesis genes. We found that cAMP-CREB mediates the expression of ASAH1 in TSC2-null cells.

A potential mechanistic insight is the appreciation the mTORi act on mTORC1 rather than mTORC2 even though the TSC1 or TSC2 mutations may affect both complexes. Logically, the mTORC2-mediated pathways are likely to contribute to any lingering phenotype of the TSC mutations. Consistent with this notion and not surprisingly, there is evidence that some of the abnormalities in TSC2-null cells are independent of mTORC1 (3). Notably, B-Raf kinase activity is reduced in TSC2-null cells due to an mTORC1-independent action of Rheb (55, 56). *Tsc1^-/-^* and *Tsc2^-/-^* MEFs have a higher percentage of cilium-containing cells compared to controls and the mTORi rapamycin has no effect on the abundance of cilia (57). Moreover, increasing evidence suggests that sphingolipid metabolism, a well-known target of mTORC2-mediated pathways may be of importance. The expression of ASAH1 is regulated via CREB-mediated transcriptional machinery. In a preliminary study, we found that TSC2-null cells accumulated higher levels of cellular cAMP relative to TSC2-addback cells, supporting a role of cAMP on viability or TSC2-null cells. cAMP regulates gene expression via the cAMP response element binding protein (CREB), a nuclear factor that is regulated by protein kinase A phosphorylation.

Renal angiomyolipomas are a major clinical manifestation of TSC. These tumors are composed of aberrant blood vessels, smooth muscle, and fat cells, representing the “mesenchymal” characteristic (45). We found increased DEGS1 and ASAH1 protein in renal angiomyolipoma cells from three patients. We have shown that TSC renal angiomyolipomas express abundant DEGS1, ASAH1 and S1PR1 proteins relative to adjacent normal kidney tissues. Treatment with rapamycin alone has a cytostatic arresting impact rather than remission-inducing effect in clinical trials and in preclinical models of TSC. These data suggest that within the context of mTORC1 inhibitor actions, there are primary pathways provoked by TSC mutations that promote persistent proliferative states. Our preliminary studies show that suppression of ASAH1 using 17a and ASAH1-shRNA attenuates xenograft tumor progression and growth of renal cystadenomas in Tsc2+/-AJ mice. We postulate that DEGS1 and ASAH1 activation leads to increased sphingolipid biosynthesis, thereby enhancing TSC tumor cell viability and growth via S1PR1 in preclinical models. Interestingly, studies have shown that *Tsc2^-/-^* MEFs are resistant to ceramide-triggered death (58). Collectively, these results suggest a cell context-specific regulation of sphingolipid metabolism and actions in TSC.

In summary, our studies identify a rapamycin-insensitive disorder of sphingolipid metabolism in TSC2-null cells and tumors and validate the novel hypothesis that TSC2 regulates sphingolipid production and action via ASAH1. Targeting aberrant sphingolipid metabolism pathways may have therapeutic value in TSC and LAM, and possibly in mTORC1-hyperactive neoplasms.

## Methods

### Cell culture and reagents

Eker rat uterine leiomyoma-derived (ELT3) cells (59, 60) were kindly provided by Dr. C Walker, Institute of Biosciences and Technology Texas A & M University, Houston, TX. *Tsc2^+/-^* A/J mice at 9 months old were obtained as a generous gift from Dr. Steve Roberds, Tuberous Sclerosis Alliance, Washington D.C. ELT3 cells stably expressing luciferase (ERL4) (20) and LAM patient-associated angiomyolipoma-derived (621-101) cells were kindly provided by Dr. EP Henske, Brigham and Women’s Hospital-Harvard Medical School, Boston, MA (20, 61). Cells were cultured in DMEM/F12 supplemented with 10% FBS, and 1% penicillin-streptomycin-amphotericin B (PSA).

### Confocal Microscopy

ELT3 cells and LAM-derived cells were plated overnight on glass coverslips in 12-well tissue culture plates. Cells were serum starved overnight, and then treated with 20 nM rapamycin for 24 hrs. Cells were rinsed with PBS twice, fixed with warm 4% paraformaldehyde, permeabilized with 0.2% Triton X-100, blocked in 3% BSA/PBS for 1 hr, and then incubated with primary antibody 1% BSA in PBS for 1 hr followed by secondary antibodies for 1 hr at room temperature. Images were captured with a FluoView FV-10i Olympus Laser Point Scanning Confocal Microscope.

### Expression array analysis

Re-analysis of previously published expression array data (GEO accession number GSE16944) (Lee et al, 2009) was performed using an online tool GEO2R. Transcript levels of were compared between TSC2-deficient (TSC2-) and TSC2-addback (TSC2+) cells, or rapamycin-treated and vehicle-treated TSC2-deficient (TSC2-) cells.

### siRNA transfections

Two separate human ASAH1-siRNAs (50 nM) (Dharmacon,) were transfected into 621-101 cells using Lipofectamine RNAiMAX (Invitrogen) according to the manufacturer’s protocols. Cells were harvested 48 hours post-transfection.

### shRNA downregulation

293T packaging cells were transfected with ASAH1 shRNA, or non-Targeting shRNA vectors using Mirus Trans-IT TKO Transfection reagent (Mirus). 621-101 cells were transduced with lentiviruses for 48 hr and then selected against puromycin. Stable clones were harvested for future experiments.

### Quantitative RT-PCR

RNA from cultured cells was isolated using RNeasy Mini Kit (Qiagen). Gene expression was quantified using One-Step qRT-PCR Kits (Invitrogen) in the Applied Biosystems Step One Plus Real-Time PCR System, and normalized to beta-actin (human) or alpha-tubulin (rat). Primers used are listed in supplementary table.

### Immunofluorescence staining

Sections were deparaffinized, incubated with primary antibody (1:100 in PBS+3%BSA) and smooth muscle actin (SMA, 1:200 in PBS+3% BSA, Santa Cruz, # sc32251), and secondary antibodies (1:1,000, Invitrogen, #A-21202 and #A10042). Images were captured with Fluorescence Microscope (Olympus BX60).

### Fluorescence-activated cell sorting

Annexin V: FITC Apoptosis Detection Kit I (BD #556547) was used to stain 621-101 cells according to the manufacturer’s protocols. FACS analysis was carried out on BD FACSCanto™ II.

### LC/MS-MS

Plasma samples were stored at −80°C and allowed to thaw at room temperature. Extraction of S1P from plasma samples (25 μL) was conducted with a modified method based on (62). Briefly, Milli-Q water (25 μL) was first added to each sample followed by addition of 50 μL of LPA solution (30 mM citric acid and 40 mM sodium phosphate, pH 4.0). After 30-second vortexing, the samples were extracted with 200 μL of extraction solution consisting of 40 ng/mL d17: S1P (IS) in MeOH. After centrifugation for 3 min at 14,000 rpm at room temperature, 200 μL of supernatant was removed and transferred into a 96-well plate for LC-MS/MS analysis. Quantification of S1P was performed by a Waters Xevo TQ-S Micro triple quadrupole mass spectrometer coupled with a Waters Acquity UPLC H-Class liquid chromatography system (Waters, Milford, MA). Multiple Reaction Monitoring (MRM) mode was used for quantification of S1P and chromatography was conducted on a Thermo BETASIL C8 column (2.1×100 mm, 5 μm, Thermo-Fisher, Waltham, MA). The optimal signal for the ion pairs of S1P was achieved in positive ion mode with the following instrument settings: capillary voltage, 3.5 kV; cone voltage 20 V; desolvation temperature, 400°C; desolvation gas flow, 800 L/h. Helium was used as the collision gas. A gradient mobile phase was used with a binary solvent system, which ramped from 50% solvent B to 100% solvent B over 1 min, after which it was held for 2.5 min, then changed to 50% solvent B over 0.1 min before being held for 1.4 min for re-equilibration. The total run time was 6 min using a flow rate at 0.6 mL/min. The column temperature was set at 60 °C and the autosampler temperature was set at 5 °C. Solvent A consisted of water/methanol/formic acid (97/2/1 v/v); solvent B consisted of methanol/acetone/water/formic acid (68/29/2/1 v/v/v/v) (63). Details of the MRM transitions monitored and the cone and collision voltages, and retention times for analytes and internal standard are shown in Table 1. Data were acquired and processed with Masslynx 4.1 software (Waters). Acetonitrile, methanol, acetone and formic acid (99%) were purchased from Fisher Science (Waltham, MA). Citric acid was purchased from Sigma-Aldrich (St Louis, MO). Sodium phosphate was purchased from Fisher Science (Waltham, MA). Sphingosine 1-phosphate (d18:1) (S1P) and sphingosine-1-phosphate (d17:1) (internal standard) were purchased from Avanti Polar Lipids (Alabaster, AL).

**Table 1.** Optimized conditions for quantification of S1P and internal standard by ESI-UPLC-MS/MS.

| Analyte | MRM transitions<br>(m/z) | Ion | Cone voltage<br>(V) | Collision<br>(V) | Retention time<br>(min) |
| --- | --- | --- | --- | --- | --- |
| S1P | 380.4 -> 264.4 | Quantifier | 20 | 5 | 2.73 |
|  | 380.4 -> 82.0 | Qualifier | 20 | 5 | 2.73 |
| d17: S1P (IS) | 366.4 -> 250.4 | Quantifier | 20 | 5 | 2.66 |

Calibration standards containing 0, 20, 50, 100, 200, 500, 1000 and 2000 ng/mL of S1P and QC samples containing 20, 60, 180 and 1800 ng/mL of S1P were prepared by pipetting the appropriate volumes of S1P working standard solutions into the charcoal stripped human serum (Golden West Diagnostics, Temecula, CA). Calibrators and QC samples were extracted using the same procedure as of the plasma samples.

### Cell viability assay

Cell viability was estimated using the 3-(4,5-dimethylthiazol-2-yl)-2,5-diphenyl tetrazolium bromide (MTT) assay. TSC2-null 621-101 and TSC2-addback 621-103 cells were plated in 96 well plates (2×10^3^ cells/well). Cells were incubated in 37°C CO_2_ incubator for 24h prior to treatment with escalating concentrations of fludarabine (0-500uM; #118218 Selleck Chemicals LLC, Tx, USA) for 72h. 25ul of MTT solution (2.5mg/ml growth media) was added to each well, followed by 4h incubation of plates at 37°C. Formation of formazan crystals was observed under the microscope. The MTT solution was replaced by 125ul of 0.04N HCl in isopropanol development solution, followed by 1.5h incubation of plates at 37°C. Cell viability was determined by measuring absorbance at 560nm with a reference of 650nm. Growth inhibitory power was analyzed using CalcuSyn Software.

### Immunohistochemistry

Sections were deparaffinized, incubated with primary antibodies and biotinylated secondary antibodies and counterstained with Gill’s Hematoxylin.

### Animal studies

Female C.B_Igh-1b/IcrTac-Prkdcscid (SCID) mice at 4-6 week of age were purchased from Taconic Biosciences (Germantown, NY) and used in this study. 2 x 10^5^ cells were injected into mice intravenously as previously described (20). Animal health was monitored daily during the tumor experiments. All mice were euthanized by carbon dioxide (CO_2_) inhalation via compressed gas after the last image was taken.

### Bioluminescent reporter imaging

Ten minutes prior to imaging, mice were given D-luciferin (120 mg/kg, i.p., PerkinElmer Inc., # 122799). Bioluminescent signals were recorded using the Xenogen IVIS Spectrum System. Total photon flux of chest regions was analyzed as previously described (20).

### Drug treatment *in vivo*

Upon tumor onset, mice were randomized and treated with the following six groups: vehicle, rapamycin (1 mg/kg/day, i.p.), 17a (10 mg/kg/day, i.p.) (64), rapamycin plus 17a (1 mg/kg/day i.p. + 10 mg/kg/day i.p.), for four weeks. Treatments were given 5 times/week for four weeks (or less if necessary, because of animal pain or distress). A separate cohort of mice inoculated with tumor cells were treated for four weeks, and then observed for tumor regrowth for eight additional weeks after discontinuation of treatment.

For 17a in *in vivo* studies, a cohort of A/J Tsc2+/- mice at 9 months old were obtained as a generous gift from the TSC Alliance. The TSC2 heterozygous model was made from deletion of exons 1-2 of the TSC2 gene on chromosome 16 as previously described. Mice were randomly divided into four treatment groups: vehicle group, rapamycin single treatment group, 17a treatment group, then rapamycin and 17a dual treatment group (N = 6-8/group). Within two hours prior to treatment, rapamycin (Enzo) were diluted with 10% PEG300+0.5% Tween 80 solution, 17a (Cayman) was diluted with 1XPBS (corning). Rapamycin was given i.p at 3mg/kg per day, and 17a were given i.p at 10mg/ml per day. Animal weight and health conditions were monitored on weekly basis. Renal tumor burdens were assessed by Magnetic Resonance Imaging (MRI) every 4 weeks.

### T2 weighted MRI to assess renal tumor progression in *Tsc2^+/-^* mice

MRI scans were carried out at the In vivo Microimaging Laboratory at Cincinnati Children’s Hospital Medical Center using a high-field (9.4 T) small bore (20 cm) Bruker Biospec 94/20 MRI/MRS spectrometer equipped with S116 high performance gradient insert and avance II electronics (66). Scans were acquired using a 72 mm id transmit/receive quadrature polarized birdcage rf coil. Mice were anaesthetized with 5% isoflurane in a 40% O_2_/air mix at 1.2 L/min, injected with 2×0.2 ml of 4% glucose/0.18% saline solution subcutaneously and transferred to specialist MRI bed model. T2 weighted, respiratory gated, fat suppressed RARE scans were performed with an FOV of 10.0×4.0 cm, amatrix of 640×256, and 64×0.5 mm coronal slices, a TEeff of 26 ms, a TR of 4100 ms, a RARE factor of 4, a BW of 100 kHz. The volumes of renal lesions and whole kidneys were measured using the software Analyze 9.0 (Analyze Direct, Inc.).

### Development of renal cystadenomas

*Tsc*2^+/-^ mice develop renal cystadenomas at high frequencies. Six treatment groups were studied, including one vehicle control group. We treated mice at five months of age with rapamycin (2 mg/kg/day, i.p.), 17a (10 mg/kg/day, i.p.), and rapamycin plus 17a (2 mg/kg/day, i.p. + 10 mg/kg/day, i.p.). We have chosen these doses according to reports in multiple preclinical studies. Treatments will be given five times/week. For *Tsc*2^+/-^ mice, eight animals were treated for four months and then sacrificed at nine months of age. To examine tumor regrowth, eight mice were withdrawn from treatment for two months and then sacrificed at 11 months of age.

### Statistical analyses

Data represent mean ± SD. Statistical analyses were performed using two-tailed Student’s t-test when comparing two groups for in vitro and in vivo studies, and one way ANOVA test (Dunnett’s multiple comparisons test when comparing multiple groups with control group, Tukey’s multiple comparisons test when making multiple pair-wise comparisons between different groups) for multiple group comparisons. A *P* value less than 0.05 was considered significant.

### Study approval

The University of Cincinnati Standing Committees on Animals approved all procedures described according to standards as set forth in The Guide for the Care and Use of Laboratory Animals. The Institutional Review Board of the University of Cincinnati approved all human relevant studies.

